# Applying rearrangement distances to enable plasmid epidemiology with pling

**DOI:** 10.1101/2024.06.12.598623

**Authors:** Daria Frolova, Leandro Lima, Leah Roberts, Leonard Bohnenkämper, Roland Wittler, Jens Stoye, Zamin Iqbal

## Abstract

Plasmids are a key vector of antibiotic resistance, but the current bioinformatics toolkit is not well suited to tracking them. The rapid structural changes seen in plasmid genomes present considerable challenges to evolutionary and epidemiological analysis. Typical approaches are either low resolution (replicon typing) or use shared k-mer content to define a genetic distance. However this distance can both overestimate plasmid relatedness by ignoring rearrangements, and underestimate by over-penalising gene gain/loss. Therefore a model is needed which captures the key components of how plasmid genomes evolve structurally – through gene/block gain or loss, and rearrangement. A secondary requirement is to prevent promiscuous transposable elements (TEs) leading to over-clustering of unrelated plasmids. We choose the “Double Cut and Join Indel” model, in which plasmids are studied at a coarse level, as a sequence of signed integers (representing genes or aligned blocks), and the distance between two plasmids is the minimum number of rearrangement events or indels needed to transform one into the other. We show how this gives much more meaningful distances between plasmids. We introduce a software workflow *pling* (https://github.com/iqbal-lab-org/pling), which uses the DCJ-Indel model, to calculate distances between plasmids and then cluster them. In our approach, we combine containment distances and DCJ-Indel distances to build a TE-aware plasmid network. We demonstrate superior performance and interpretability to other plasmid clustering tools on the “Russian Doll” dataset and a hospital transmission dataset.

**Impact statement:** Studying plasmid transmission is a necessary component of understanding antibiotic resistance spread, but identifying recently related plasmids is difficult and often requires manual curation. Pling simplifies this by leveraging a combination of containment distances and rearrangement distances to cluster plasmids. The outcome are clusters of recently related plasmids with a clear backbone and relatively large core genomes, in contrast to other tools which sometimes overcluster. Additionally the network constructed by pling provides a framework with which to spot evolutionary events, such as potential fusions of plasmids and spread of transposable elements.

**Data summary:** Supplementary information and figures are available as an additional PDF.

The tool presented in this paper is available under https://github.com/iqbal-lab-org/pling. Additional computational analysis and scripts are described and provided under https://github.com/babayagaofficial/pling_paper_analyses. The sequence data used can be found under BioProject no. PRJNA246471 in the National Center for Biotechnology Information for the “Russian doll” dataset (https://www.ncbi.nlm.nih.gov/bioproject/PRJNA246471), and under Project no.

PRJEB31034 in European Nucleotide Archive for the “Addenbrookes” dataset (https://www.ebi.ac.uk/ena/browser/view/PRJEB30134). All other genome sequences used were sourced from PLSDB (https://ccb-microbe.cs.uni-saarland.de/plsdb/), and lists of accession numbers can be found in the additional analysis github.

## Introduction

The genetic content of a bacterial cell comprises both the chromosome, and a collection of between 1 and 10 plasmids – autonomously replicating, (mostly) circular DNA molecules, each of which is independently maintained at some copy number between 1 and around 100 [1, 2]. The chromosome is passed on directly to daughter cells, and its evolution can be modelled with phylogenetic approaches. However, the plasmids can, either independently or in collaboration, move to another potentially completely unrelated cell by various means [3]. Thus plasmids can be viewed from the perspective of a cellular lineage and its associated chromosome as extra genetic content providing further functionality, or as independent Darwinian elements moving between different environments (i.e. hosts, or even species). Plasmid genomes seem to undergo structural changes (gain/loss of genes, rearrangements) at comparable rates to mutations (although rates are hard to measure in natural populations if horizontal gene transfer is involved) [4]. Even more, unrelated plasmids within a cell can undergo recombination mediated by repeat elements, or even cointegrate to form a new combined plasmid [5–7]. In these circumstances the notion of plasmid “identity” has to be rather fluid.

The question of plasmid relatedness naturally arises in epidemiological studies, which must analyse a set of bacterial (chromosome and plasmid) genomes and determine whether there is a clonal spread, or a plasmid spread, or neither. Coarse approximations, such as classifying plasmids based on specific alleles of replicons (replication and partition genes) [8, 9] or of relaxase genes [10], are insufficiently fine-grained – sometimes the “types” can correspond to stable “backbones” (e.g. IncW [11, 12], IncP-1) but others consist of highly mosaic recombinants with no common structure (e.g. IncF [13–15]), and there can also be recombination between types [6]. Furthermore, many plasmids are not type-able at all based on existing databases. In cellular organisms (esp. bacteria), a combination of ecology and preferential recombination with genetically similar organisms leads to a natural clustering of genomes into species.

Therefore two alternative methods have been used more recently to look for natural separation into groups of plasmids. “Sketching” methods compare the k-mer (DNA word of length k) content of two plasmids, and are very effective at finding almost identical plasmids [16–18]. Alternatively, complete alignment of a pair of plasmids can be used to measure simple containment (the percentage of a plasmid that lies in an alignment block with another plasmid) [19]; this can be approximated using containment sketch approaches [20, 21]. Neither approach attempts to measure the number of genetic events separating two plasmids, and can significantly over- and under-estimate genetic distance as a result. This highlights the key challenge in detecting recent ancestry, and potentially transmission, of plasmids: there is no intuitive genetic distance which counts the number of structural events between plasmids.

We address this issue, using very recent methods from the field of combinatoric analysis of genome rearrangements to count the minimum number of (specific types of) rearrangements between two plasmids. Algorithms for counting the number of structural changes between genomes encoded as ordered lists of markers (e.g. genes) have been studied since the early work of Sankoff and Pevzner [22, 23]; see [24] for a thorough history. The key development for our needs was the introduction of the Double Cut and Join model [25] in 2005. This introduced a single operation (cut the genome in two places, creating four ends, and reconnect them in a different way) which was general enough to generate inversions, formation and reabsorption of circular intermediates, and translocations (See Figure 1).

**Figure 1:**
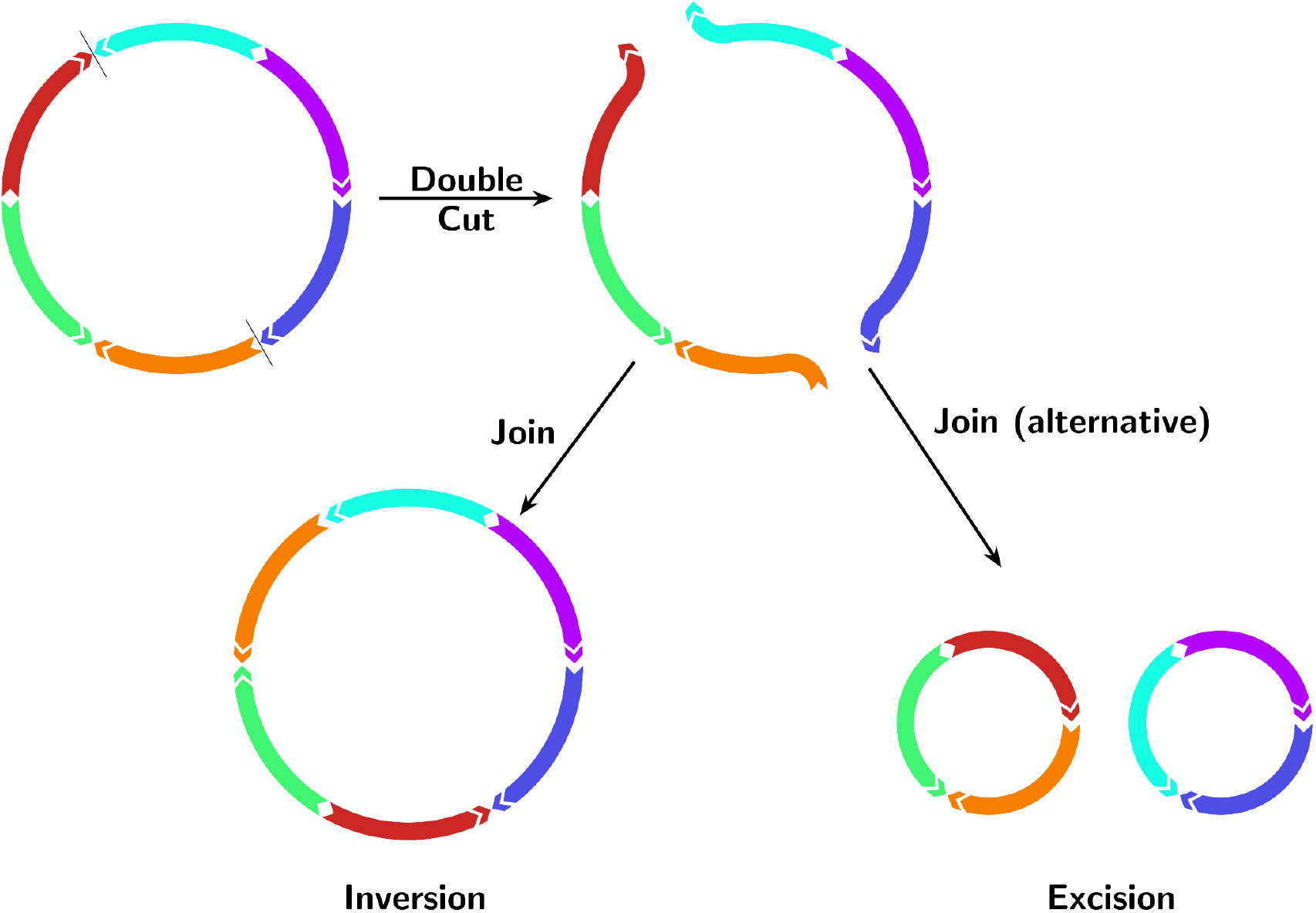
Visualisation of how the DCJ (Double Cut and Join) operation affects a circular plasmid. First there is a “double cut”, cutting the genome in two places and creating four ends. Then two pairs of ends are reconnected; if we exclude the original pairing, this leaves two possible ways to reconnect the free ends, resulting in either an excision or inversion.

These are all key “operations” for modelling plasmid evolution. The model was initially developed for genomes with identical marker content, followed by extensions that allowed for indels [26, 27], or included duplicated markers [28]. Finally, both indels and duplicates were included in an integer linear programming formulation of the problem, and implemented in the tool ding [29] in 2021. This sufficiently broadened the model that it could be effectively applied to calculate rearrangement distances between plasmids.

We develop a software workflow, *pling*, which takes a set of plasmids (as fasta files) and calculates the minimum number of DCJ, insertion or deletion events between all pairs using an updated version of ding. We demonstrate how pling can be used to infer different types of relationships between plasmids, including cointegration and transposable element (TE) spread. Furthermore, we apply pling to hospital plasmid datasets to study their epidemiology. We discover that pling is able to avoid the pitfalls of other clustering tools by accounting for structural changes with the DCJ-Indel distance, and as a result produces intuitive and easily interpretable clusters.

## Theory and implementation

### Applying DCJ-Indel to pairwise distances between plasmids

The DCJ model regards the genome as an ordered sequence of signed integers, in which each integer represents a region (commonly genes or synteny blocks), and the sign implies orientation. Multiple copies of a gene (repeats of a region) would use the same integer. The basis of the model is the double cut and join operation, in which a genome is cut at two different positions, creating four open ends, and these are joined in a new way (see Figure 1). Almost all the key operations we require in order to model structural changes of plasmids are encompassed by this model: inversions, translocations, fusions and excisions. In this paper we use the DCJ-Indel model, an extension of the classical DCJ model [25] that handles insertions and deletions. The DCJ-Indel distance between two genomes is then defined as the minimal number of DCJs, insertions and deletions necessary to transform one genome into the other [26]. As a result, the distance is a representation of how many structural changes differentiate a pair of plasmids. Bohnenkämper *et al*. developed an integer linear programming software implementation, ding, for calculating the DCJ-Indel distance between two genomes which had already been represented as integers [29]. We describe below how we built a plasmid workflow around version 2 of ding (https://gitlab.ub.uni-bielefeld.de/gi/dingiiofficial.git).

### Pling: software workflow to calculate DCJ-Indel and cluster plasmids

Pling is a software workflow designed to enable rapid calculation of DCJ-Indel distances between plasmids, construct a (TE-aware) relatedness network and then find clusters of closely related plasmids using a network community algorithm. Pling works with either synteny blocks defined at the sequence level (using nucmer [30], run by pling), or using genes defined by annotation (using bakta [31] and panaroo [32], run by pling). Since gene annotation software can underannotate plasmids and sometimes fails on diverse sets of plasmids, we find sequence-based analysis a much more reliable and rapid option and describe this in detail here. A brief summary of the annotation workflow is given in the Supplementary Material for completeness sake.

The workflow proceeds through the following steps:

*Input data*: We start from complete, circular plasmid genome assemblies.

#### Step 0: Preliminary technical issue

Technically speaking, one plasmid can always be transformed to another by two operations: one deletion removes all integers from the first plasmid, and then one insertion adds all the integers from the second plasmid. This is generally prevented by the DCJ-Indel model by only allowing deletions of overrepresented segments and insertions of underrepresented segments. Nevertheless, this phenomenon still applies in the case of two unrelated plasmids (no integer in common). Therefore to prevent such occurrences, we only calculate DCJ-Indel for pairs of plasmids which share sequence – hence the reason for the initial construction of a containment network (details below).

#### Step 1: Integerisation and construction of a containment network

The first step is to construct a network connecting all pairs of plasmids which may conceivably be related, by looking for some level of shared sequence. The measure used here is a containment distance, which is defined as follows: Say two plasmids *p*_1_ and *p*_2_ align into *N* matches *m*_*i*_ (*i* = 1,..., *N*), let *len*_*j*_ (*m*_*i*_) be the length of the match *m*_*i*_ on plasmid *p*_*j*_ and *len*(*p*_*j*_) be the length of the plasmid *p*_*j*_ (*j* = 1, 2), then if *len*(*p*_1_) < *len*(*p*_2_), the containment distance *CD* is

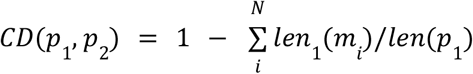

– i.e. measuring the proportion of the smaller plasmid that is not contained in the larger. This is similar to the distance used by MobMess [19], and is approximated by the containment measure used by sourmash [21]. We refer to this as the *containment network*, but note this does *not* imply that plasmids connected by an edge necessarily have one plasmid completely contained in the other – the containment terminology is inherited from other papers in the field [19].

For each pair of plasmids, we perform *integerisation* – this is the process of transforming the DNA sequences into ordered signed integer sequences. Each integer in the sequence represents a continuous block of sequence; the sign represents the orientation. We run MUMmer 3 [30] (for subprograms and parameters, refer to Supplementary Information), which gives us a set of matches between the pair of plasmids. We use these to identify synteny blocks and to assign the same integer label to them (details below). Any regions of at least 200 bases not covered by a synteny block are regarded as indels, and also assigned a unique integer. We also calculate the containment distances from these matches.

We then build a network where each node represents a plasmid, and plasmids are joined by an edge if their containment distance is below a prespecified threshold (default 0.5, or for transmission analysis 0.3). Each connected component in this network defines a plasmid community.

#### Step 2: Identify hubs

The next step is to identify *hub* plasmids, which are densely connected plasmids, whose neighbours are sparsely connected amongst each other. We find hubs by first checking if the degree of a node in the network is higher than a prespecified hub connectivity threshold (default 10), and then check if the edge density of that node’s neighbours is lower than a prespecified threshold (default 0.2), in which edge density is defined as the number of edges between nodes relative to the number of edges if all nodes were connected. Hubs are found iteratively – first all hubs are found on the original containment network, then all hubs on the network induced by removing all hubs, and so on until no more hubs are found. These hubs are typically relatively small and dominated by a transposable element, i.e. a genetic element that can insert itself at different positions, including insertion sequences and transposons [33, 34].

#### Step 3: DCJ-Indel calculation and network construction

We next calculate the DCJ-Indel distance for all pairs of plasmids that meet the prespecified containment threshold, using the software ding. We use the distances to induce a subnetwork on the containment network: any edge between plasmids whose DCJ-Indel distance is greater than a prespecified threshold (default 4) is removed. Additionally, we disconnect the network at the previously found hubs (all edges from hub plasmids are deleted), allowing the network to split into components which previously were only connected via the hubs. At this stage, any two plasmids connected by an edge share sequence, and are separated by at most 4 structural events.

#### Step 4: Network community detection

Finally, on this subnetwork we run the asynchronous label propagation community detection algorithm [35, 36] to define plasmid subcommunities. This algorithm has the benefit of not requiring prior knowledge of the number of subcommunities, and is reasonably fast.

##### Inferring integerisation from alignment: details

In this section, a *match* refers to a block of aligned sequence found by a pairwise alignment using MUMmer subprogram nucmer. If there are no duplicate or overlapping matches (i.e. each match occurs precisely once on each genome), integerisation is trivial. Assign a unique integer to each match, and then assign integers to the gaps between matches on the two genomes. However, there is a long list of potential edge cases once a block in one genome has two matches on the other, or when matches overlap (see Supplementary Figure 8 for an example). Therefore we preprocess the nucmer matches to resolve any overlaps where we can, before assigning integers. Suppose we are processing genome A and genome B. The overlap-fixing algorithm implemented in pling begins by sorting the matches according to ascending coordinates in genome A. After sorting, it iterates through every match (i.e. moves left to right on genome A), until it finds a pair of overlapping matches, upon which it:

1. Finds the coordinates of the overlap on genome B, which will result in two sets of overlap coordinates
2. Identifies which matches contain the overlap coordinates on genome B
3. Splits each of these matches around the overlap coordinates

and then continues onwards. This process is then repeated for overlaps on genome B. We assume that after the processing there are no overlaps, unless matches represent duplicate regions (and therefore share the exact same coordinates on either the reference or query genome). Then non-duplicate matches receive a unique integer, as in the trivial case, and matches representing duplicate regions all receive the same integer, and finally gaps are assigned integers. A visualisation of this full process can be found under Supplementary Figure 9.

The overlap fixing algorithm necessitates having reliable projection between reference and query coordinates on a match, which accounts for small indels within a match. This is mostly available with the help of MUMmer subprogram show-snps output, which is a list of SNPs and small indels. However when the structure of the plasmids becomes very complicated (e.g. lots of repeats offset only by a couple of bases), it becomes impossible to accurately identify which small indels belong to which match, and therefore the projection between coordinates becomes unreliable. We give details of how this is handled in Supplementary Information.

##### Software implementation details

Step 1 implicitly relies on running nucmer on all pairs of plasmids, which is the most time consuming step in our workflow. However there is no point doing this on plasmids which are not going to fulfil the containment criterion, making this step amenable to optimisation. We therefore run sourmash’s containment estimate on all pairs, which is very fast and based on k-mer sketches, to rule out pairs of plasmids which are definitely not going to be connected in the network. We rule out pairs of plasmids at a lower threshold (default 0.15) than the containment network threshold, as sourmash tends to underestimate containment and we want to avoid ruling out viable pairs. This reduces the runtime considerably. For example, on 1500 randomly selected plasmids from PLSDB [37], without sourmash prefiltering pling’s runtime was about 104 minutes, while with prefiltering it was a little under 7 minutes. We see a reduction in runtime for less diverse plasmids as well – on a set of 1500 IncF plasmids from PLSDB, without sourmash the runtime was about 273 minutes, while with sourmash it was about 79 minutes. Adding sourmash filtering also makes running smaller datasets on a laptop tenable, as the runtime for 300 IncF plasmids on a laptop with 8 cores was about 22 minutes. See Supplementary Information and Supplementary Table 1 for further details.

By default, ding relies upon the integer linear programming (ILP) library Gurobi [38], which is closed source and requires the user to sign up for a licence (free for academic use). We found an alternative library to use, GLPK [39], which is open source. As ding only supports input in the form of a Gurobi solutions file, we wrote a script to convert GLPK solution files to Gurobi solution files, which allowed us to implement GLPK as the default ILP solver for pling. We still run most of our analysis with Gurobi though, as it offers a runtime benefit: for the same 1500 IncF plasmids as above, runtime with sourmash prefiltering and GLPK was about 114 minutes, which is slower than with Gurobi by about 35 minutes. For smaller datasets the benefit is less pronounced though, as the 300 IncF plasmids completed in 24 minutes, i.e. 2 minutes slower.

## Results

We implemented a software workflow pling, to infer a relatedness network between a set of (circularised) complete plasmid genomes based on the DCJ-Indel distance. Small transposable elements such as transposons and IS elements frequently jump between unrelated plasmids, creating a false signal of relatedness for metrics which look for shared sequence. We therefore specifically design our workflow to identify “hub” nodes in the network which are hyperconnected and join otherwise unrelated plasmids, as they are potential TEs. We prevent these from connecting unrelated plasmid subcommunities (“types”) – see Methods for details of the workflow. The key aspects are: we create a containment network (edges connecting plasmids where most of the smaller is contained in the larger) to identify plasmids which share some sequence, and then a subnetwork by removing edges between plasmids separated by more than some threshold (we use 4) number of structural events (illustrated in Figure 2). By combining these three steps: sequence sharing, hub removal, rearrangement distance threshold, we arrive at a very interpretable graph which reflects what we intuitively expect in real biological scenarios.

**Figure 2.**
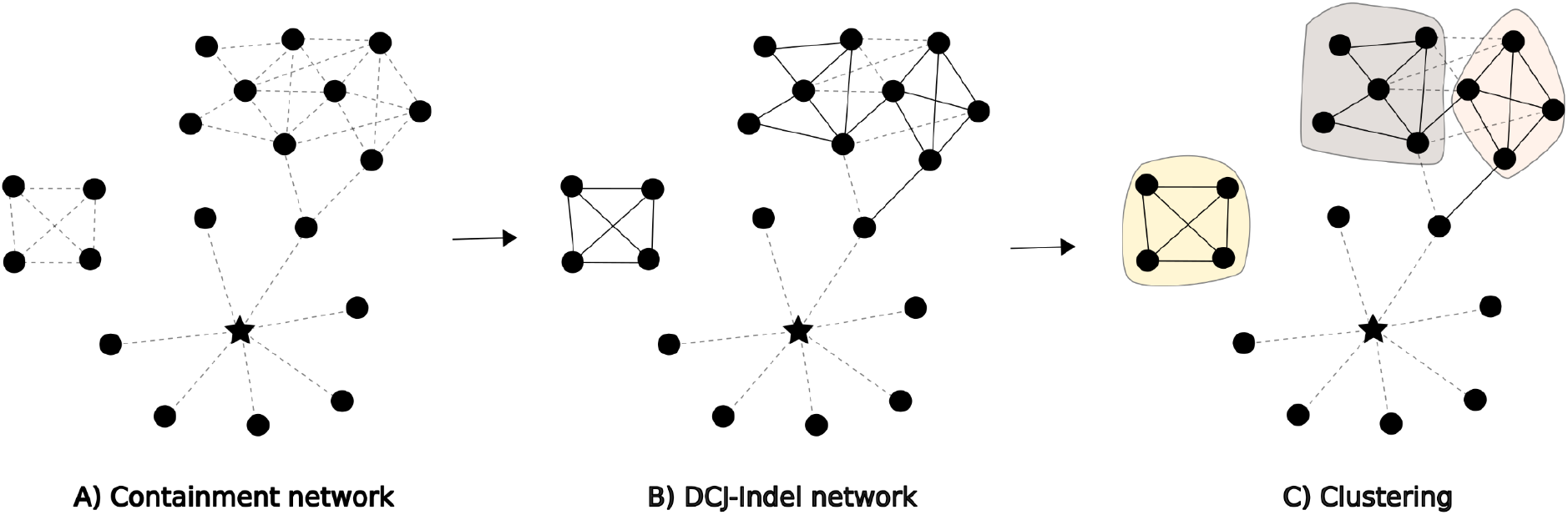
The pling workflow. A) Plasmid containment network, dotted line between nodes (plasmids) whose containment distance is less than 0.5 (if looking for broad relatedness) or \0.3 (if looking for potential transmissions). Hub node (plasmid potentially containing a transposable element) is marked by a star. B) Dotted edges separating plasmids with DCJ-Indel distance≤4 are changed to solid edges. For edges connected to a hub plasmid, even if DCJ-Indel distance≤4, the edges remain dotted. C) Asynchronous label propagation network community algorithm is used to identify plasmid subcommunities (shaded backgrounds).

### Illustrating key scenarios with the Plasmid Spectrum Dataset (PSD)

We begin by elucidating some of the different types of relationships possible between plasmids, and how these are reflected in terms of Jaccard distance (non-shared k-mers over union, estimated by mash [40]), containment distance (as implemented in sourmash [21]), and DCJ-Indel distance (pling). In order to do this, we constructed the Plasmid Spectrum Dataset (PSD), which is a dataset of plasmids we have curated from various sources to illustrate four possible scenarios of plasmid relatedness:

A. High sequence similarity, and few rearrangements or indels
B. High sequence similarity, and many rearrangements or indels
C. Fusion of plasmids
D. Promiscuous TE spread across many divergent plasmids

In Figure 3, we show how these different genetic distances capture (and fail to capture) these differences. In Figure 3A) we see the simplest case, almost a “positive control” – two plasmids which are identical except for a relatively small indel – here all genetic distances are low. Unfortunately low Jaccard or containment distances are not sufficient to guarantee two plasmids are similar; in Figure 3B) we see two evolutionarily diverged plasmids, separated by many structural changes, but nevertheless sharing many k-mers, which confounds both Jaccard and containment. Only DCJ-Indel identifies that there are many structural changes (DCJ-Indel distance 88). Another common occurrence is for one plasmid to be mostly contained in another. In Figure 3C), we show such a pair. Since Jaccard distances favour genomes of similar sizes, the distance for this pair is quite high, whereas the containment distance behaves correctly, with distance zero. This does highlight that containment distance of zero does not imply plasmids are identical. The DCJ-Indel distance of 1 is also low, since it counts that the fusion event can be done in one DCJ operation, thus quantifying the number of genetic events apart in a way the other measures do not. Finally, in Figure 3D) we show an example which confounds all methods: a set of unrelated plasmids except for the fact that they all contain a Tn4401 transposon. All of the distances are low between this plasmid and the rest of the plasmids in this example. Plasmid relationships like these confuse clustering algorithms due to having a single highly connected plasmid, which can then falsely pull together divergent plasmids into a singular cluster.

**Figure 3.**
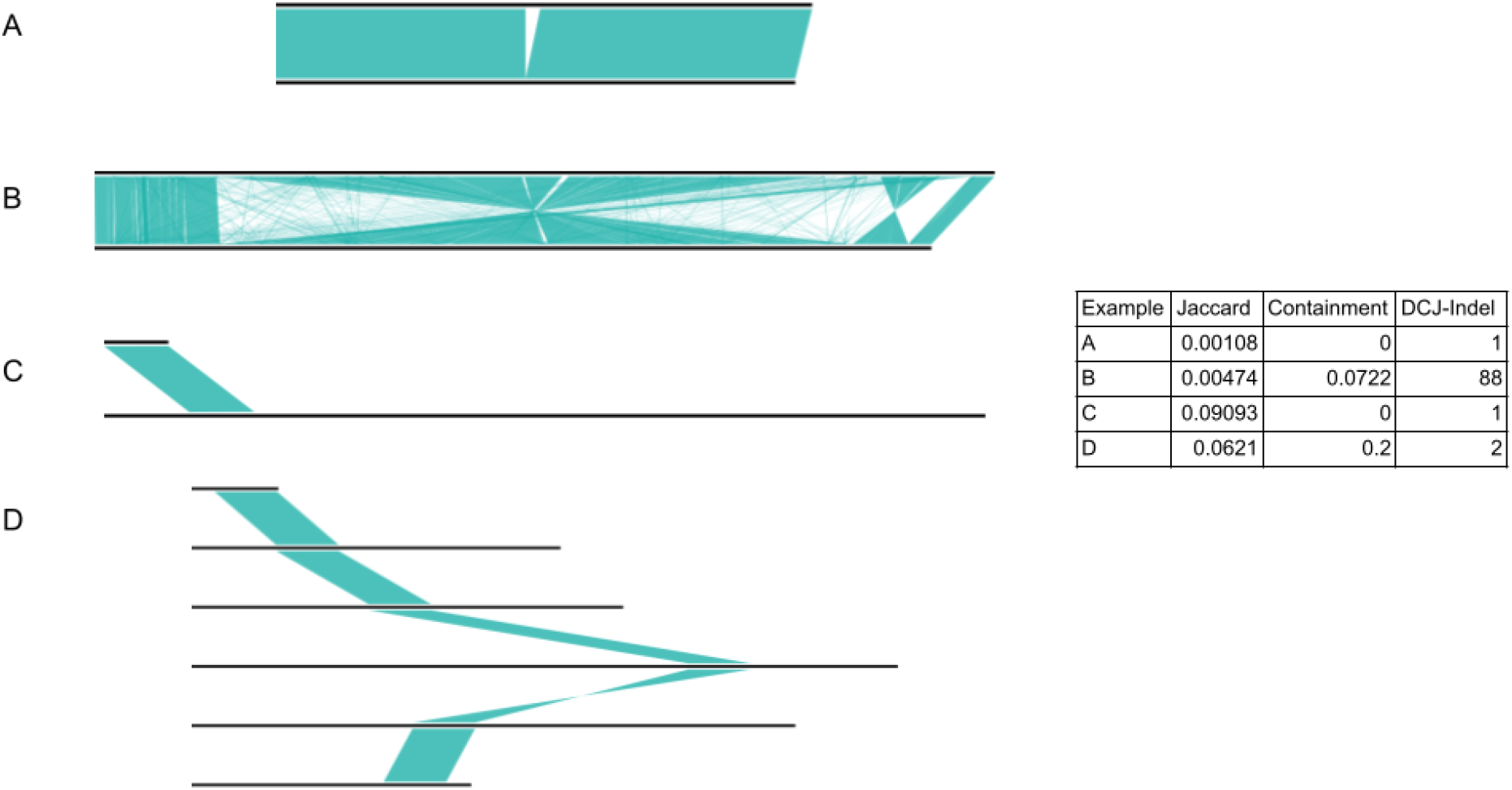
Impact of typical plasmid genetic variation on different genetic distance metrics. We show Jaccard distance (non-shared k-mers over union, calculated with mash), Containment distance (k-mers unique to smaller plasmid over k-mers in smaller plasmid, calculated with sourmash), and DCJ-indel distance, calculated with pling, for the Plasmid Spectrum Dataset. A)Two plasmids that differ only by a small indel: all distances are low; only DCJ-Indel reflects that the distance is just 1 evolutionary event. B)Two plasmids that are very diverged through many structural events, but share many k-mers: only DCJ-Indel captures the structural divergence – 88 genetic events apart. C)One plasmid contained in another: all the distances are low. Relatively high Jaccard distance due to very different genome sizes; low containment and DCJ-Indel distances as expected. Cannot identify that one is engulfed in the other without also looking at lengths. D)Several unrelated plasmids which only share a transposon. In the table the median distance of all the plasmids to the topmost plasmid is given (for the distances per pair, see Supplementary Table 2). None of the distances are particularly high – this seems a case where the distances all fail.

Workflows based on sequence similarity (Jaccard and containment) generally try to mediate this issue through thresholding, which causes a tradeoff between needing the similarity threshold low to not separate plasmids with large (but few) structural changes, and needing it high to avoid confusion from TEs.

### Combining multiple distance measures and network structure to reduce TE noise

Pling sidesteps the trade-off described above by clustering with both containment and DCJ-Indel distances, and identifying hub plasmids which connect many otherwise-unrelated plasmids (Methods). This lets us keep containment distance thresholds low so we do not miss any related plasmids, and then remove any false connections between plasmids by filtering by DCJ-Indel distance. This does not guarantee avoiding TE problems; but constructing the containment network can give enough context to differentiate between pairs with a genuine recent relationship (example C), and divergent pairs connected through TE promiscuity (example D).

### Benchmarking pling on a hospital dataset with nested Russian doll-like structural variation

We selected a small dataset from the famous “Russian doll” paper from Sheppard *et al*. [41] to compare pling to other clustering approaches. In this paper carbapenem resistant isolates containing the carbapenemase gene, *bla*_KPC,_ from a single hospital outbreak were studied, and found to be related by a hierarchy of MGE movement, including transposons and plasmids. Plasmids from 17 randomly selected *Enterobacteriaceae* isolates from the outbreak were studied in detail. All the resistant plasmids were categorised into clusters using manual curation based on BLAST, but only for one cluster were similar non-resistant plasmids identified. We clustered all plasmids >5kb found in the isolates (N=62), and compared results with MOB-suite [16] and mge-cluster [17], as both of these tools are, like pling, based on clustering with a genetic distance. Since the data were collected from August 2007 to December 2012, and we wanted to be able to capture plasmid changes, we first used the broader “primary clustering” in MOB-Suite, and later followed up with the secondary clustering which looks for very closely related plasmids. We do not look at typing from PTUs [18] in depth, as none of the plasmids belong to known PTUs.

Firstly, we immediately recover the clustering of the subset of plasmids that Sheppard *et al* analysed in detail (orange in Figure 4a,b); this cluster is a well-known example of why clustering plasmids is challenging because one plasmid is more than double in size than the others. Resistant plasmids that were originally clustered together are in the same pling cluster, and singleton plasmids are also the same as in Sheppard *et al*. We also find some novel clusters formed of non-resistant plasmids, and extend Sheppard’s clusters with non-resistant plasmids.

**Figure 4.**
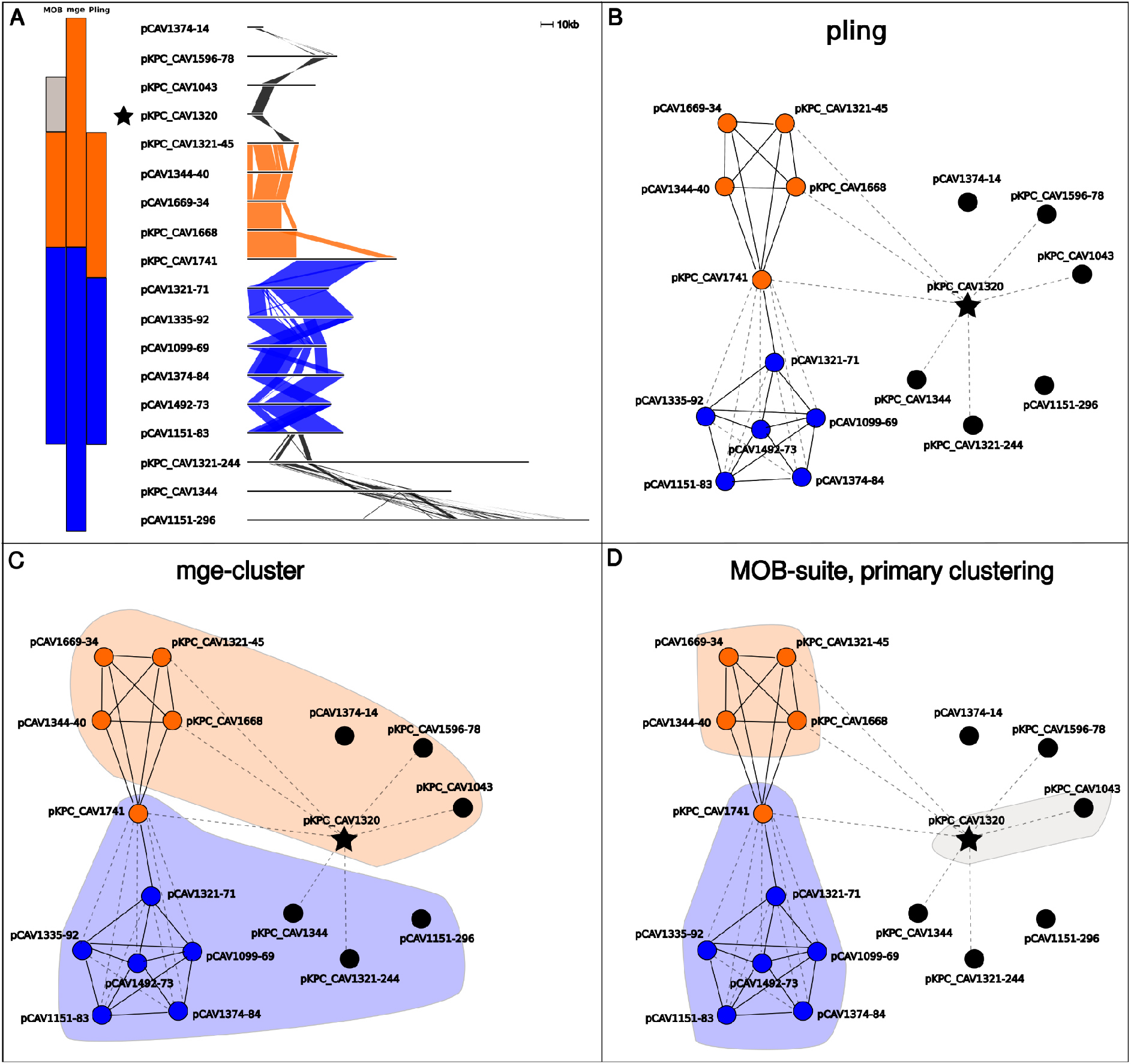
Comparing clustering on a subset of Russian dolls data. A) An alignment of a subset of the Russian Doll plasmids. The coloured bars to the left of the alignment signify to which cluster (subcommunity) a plasmid was assigned, with each column representing a tool and each colour representing a cluster (left to right: MOB-suite, mge-cluster, pling). In the alignment, shaded areas indicate shared sequence between adjacent plasmids, coloured by pling clustering assignment. The star indicates a hub plasmid. B) The pling network of the plasmids in the alignment, in which each node represents a plasmid, dashed lines represent that containment distance≤0.5, solid lines represent that DCJ-Indel distance≤4, and the star denotes a hub plasmid. The hub plasmid contains the Tn4401 transposon which was found across many of the resistant plasmids in Sheppard *et al*, and is exactly the black region in the alignment of the hub in A. Orange nodes all belong to one cluster (subcommunity), as do blue nodes, while black nodes are all singletons. C), D) Again the pling network, but with shaded areas added to show how mge-cluster (C) and MOB-suite (D) cluster these plasmids. The shaded colours correspond to those on the bars in the upper left panel.

Mge-cluster is not concordant with the Sheppard’s clusters, and the differences tend to arise from assigning singletons to clusters. It appears that mge-cluster is prone to being confused by hub plasmids, which leads to inappropriate clustering of many divergent plasmids together, as in Figures 4 and 5. MOB-suite’s primary clustering is also affected by hub plasmids, as the example in Figure 4 is exactly the same as in Figure 3D discussed above, but MOB-suite still pairs it with another singleton plasmid. Typically hub plasmids are relatively small plasmids which consist mostly of a highly mobile region found in diverse plasmids. Since pling identifies and isolates hub plasmids before clustering, we avoid this issue and have congruence with Sheppard’s clusters.

**Figure 5.**
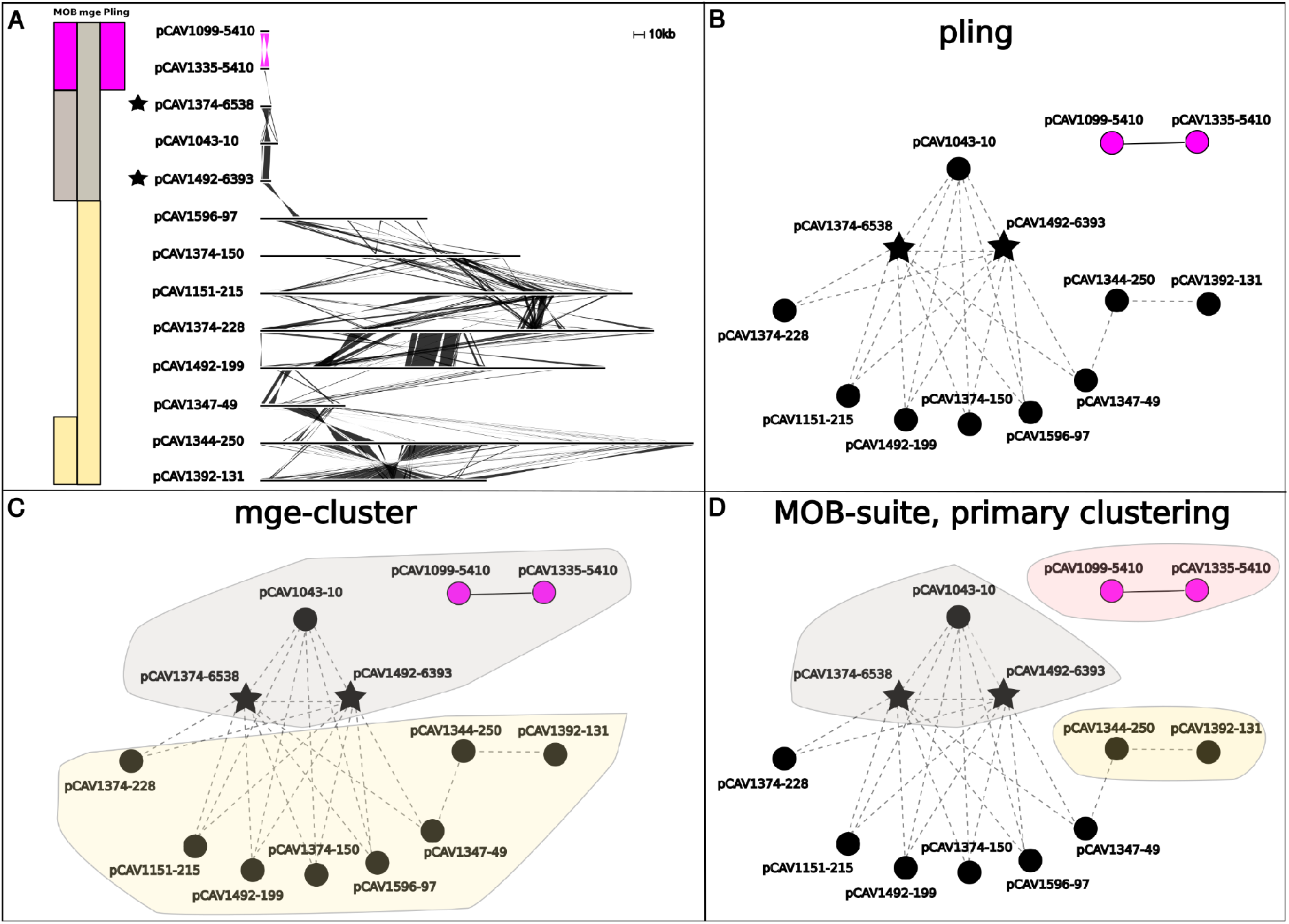
Russian dolls subset with two hub plasmids and a high DCJ-Indel pair. A) An alignment of a subset of the Russian Doll plasmids. The coloured bars to the left of the alignment signify to which cluster a plasmid was assigned, with each column representing a tool and each colour representing a cluster (left to right: MOB-suite, mge-cluster, pling). In the alignment, pink and black areas indicate shared sequence between adjacent plasmids, coloured by pling clustering assignment. B) The pling network of the plasmids in the alignment, in which each node represents a plasmid, dashed lines represent that containment distance≤0.5, solid lines represent that DCJ-Indel distance≤4, and the stars denote hub plasmids.The two pink nodes are a pair, while black nodes are all singletons. C), D) Again the pling network, but with shaded areas added to show how mge-cluster (C) and MOB-suite (D) cluster these plasmids. The shaded colours correspond to those on the bars in the upper left panel.

The dataset also contains an example similar to that in Figure 3B – there are two large plasmids (pCAV1344-250, pCAV1392-131) with very high sequence similarity, but very high DCJ-Indel distance (=40), also easily visible in their alignment in Figure 5.

Mge-cluster and MOB-suite also disagree with the assignment of plasmid pKPC_CAV1741 in Figure 4. Both tools split off the plasmid from the orange cluster and join it with plasmids that form the blue cluster in pling’s subcommunities. The plasmid is a cointegrate, and the orange and blue clusters both contain a distinct parent of it. Hence mge-cluster’s and MOB-suite’s assignment of the cointegrate is not unreasonable, as it groups it with its blue cluster parent. However, plasmids in the blue cluster generally had higher DCJ-Indel distances to the cointegrate plasmid, than the other orange cluster plasmids, supporting pling’s decision. Finally, the natural follow-up with MOB-suite is to try the secondary clustering, which returned almost identical results to pling, disagreeing only on the co-integrate parent. Thus this corrects the major error in MOB-suite primary clustering, where pCAV1344-250 and pCAV1392-131 had been clustered together despite their structural divergence.

### Understanding epidemiology of Addenbrookes hospital plasmids

We studied pling’s utility for identifying transmission from a dataset of plasmids from 72 carbapenemase-producing bacteria isolates (N=193), from a paper studying plasmid movement in carbapenem resistant isolates sampled over the course of 6 years from Addenbrookes hospital in Cambridge, UK [42]. We took the plasmids from that study, and clustered them with a relaxed mash distance threshold of 0.01, and then used alignments and additional metadata to manually curate these clusters, creating a “curated truth” of 26 clusters and 105 singletons. We compare these clusters to the pling clusters created with a containment threshold of 0.3 and DCJ-Indel threshold of 4. We changed the containment threshold from the default of 0.5 to a stricter value of 0.3 to ensure that only plasmid clusters with recent evolutionary history were identified. We compared with mge-cluster (default parameters) and MOB-suite (secondary clustering, intended for identifying transmissions).

We chose to compare clustering with the metric introduced by van Dongen [43–45], which counts how many splits and joins are necessary to transform one clustering into another. Pling identified 25 clusters, 19 of which were identical to truth clusters. The truth consisted of 105 singletons, and 88 plasmids associated with a cluster. Pling identified 105 singletons, and assigned 88 to a cluster, where 12 plasmids would have to be reassigned to get the truth clustering (∼6% of plasmids). MOB-suite and mge-cluster had less agreement with the truth, recapitulating fewer clusters and singletons (would have to reassign ∼12% and ∼42% of plasmids respectively); see Supplementary Figure 2. Furthermore we found that pling tended to produce clusters with larger relative core (median of core size as a percentage of genome size) than mge-cluster and MOB-suite, even for larger clusters, as can be seen in Supplementary Figure 1. This can make incorporating typical SNP-based analysis methods more likely to succeed for plasmids as there is more backbone available for SNPs to occur on.

We examined where pling’s disagreements with the truth came from, to establish if these were genuine misassignments, or discovery of novel relationships within the dataset. Disagreements from the truth came from either splitting plasmids from truth clusters, merging truth clusters (clusters 25 and 123, total of 5 plasmids) or adding singletons to clusters (one added to truth cluster 16, one novel cluster of 3 plasmids, and two novel pairs). There were four examples of merging clusters and adding singletons to clusters, where alignment supports pling’s choice over the original “truth” (Supplementary Figures 3-7). In the first of these four, two clusters (25 and 123) are merged (Supplementary Figure 3) revealing that the same plasmid is prevalent across ST2 of *Acinetobacter baumannii* (present in 4/5 ST2 isolates), and also shared in ST664.

In the second case, pling’s addition of a singleton plasmid to truth cluster 16 revealed sharing between *Citrobacter freundii* and several other species. Figure 6 shows the spread of plasmids shared between *Klebsiella pneumoniae* and *Escherichia coli*, with this plasmid cluster being denoted by an orange dot. We can see that there is both intraspecies and interspecies transmission of this plasmid cluster, as it is found on two different branches of *Klebsiella pneumoniae* tree, and on the *E. coli* tree. There are two plasmids from this cluster missing in this figure, as one is in an *Klebsiella aerogenes* isolate, while the other is in *Citrobacter freundii*, i.e. the singleton mentioned above.

**Figure 6:**
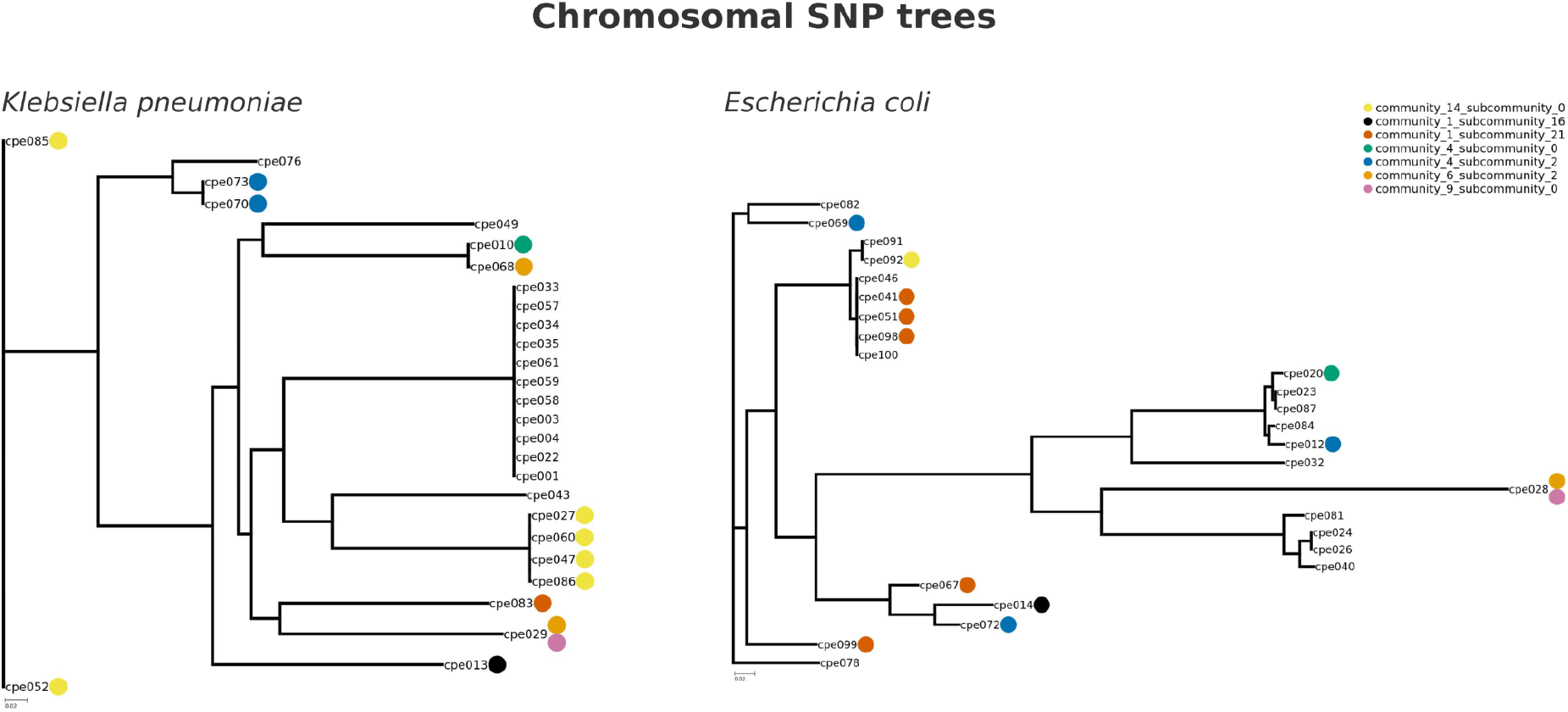
SNP trees of *Klebsiella pneumoniae* (left) and *Escherichia coli* (right) isolates from Addenbrookes hospital. The dots beside isolate names indicate the presence of a plasmid from a specific cluster, with one colour assigned to a cluster. Only the presence of plasmids which are shared across both *Escherichia coli* and *Klebsiella pneumoniae* is shown here.

For the remaining discordances, we could not come to strong conclusions; when constructing our truth set, we found no single assembler would reliably assemble and circularise the chromosome and plasmids for all samples. Therefore we chose the best assembly for each sample, but it remains likely that there are some assembly errors within the dataset, which would inflate DCJ distance.

## Discussion

It is worthwhile to view the work from this study in a wider context. Each bacterial cell was born through binary fission of a parent, leading to a cellular genealogy back in time. Bacterial DNA mostly stays confined to its own cells, and mutations happen every few generations, so analysis of contemporary genomes reveal mutational patterns from which we can infer a phylogeny, a genetic analogue of the cellular genealogy. We have a mature understanding of how recombination and horizontal gene transfer interfere with this tree-based model of evolution, and all-against-all comparisons show bacterial genomes naturally group in clusters with Approximate Nucleotide Identity (ANI) >95%, which coincide with our species definitions. Species exist as a phenomenon for bacteria – the cells are closely related in a cellular family tree, the genomes are closely related in a core-genome phylogeny, and we can recover species clusters via ANI. None of this holds for plasmids: they have no cellular identity but exist only at the DNA level within different bacterial host species, sharing those cells with other unrelated plasmids. Plasmid genomes can change structurally as or more often than mutations occur – inversions, gene gain and loss causing huge size changes, and most dramatically, fusion of plasmids within the same cell, mediated by various mechanisms. Naive clustering using Jaccard or containment metrics estimated with a subsampled set of k-mers (e.g. mash, sourmash) is only partially effective for three reasons: first, and fundamentally, the edge weights are based on k-mers, not genetic events; second, as shown in Figure 3, closeness in Jaccard sense does not necessarily imply closely related for plasmids; third, plasmids with TEs which connect multiple unrelated plasmids can still be pretty big, so simple size filters will not work. It therefore is very hard to tell if there are natural groupings of plasmids using these tools.

Due to the challenges described above, it is apparent that a method to evaluate structural genetic events is critical to understanding plasmids. While there has long existed a wealth of rearrangement distances that do exactly this, they have seen limited application in practice. This is partially because most of these models have assumptions on the genome that are too limiting for real genomes, but our model of choice, DCJ-Indel, has had recent developments that solve this issue (ding) [29]. The combination of this recent work, and the fact that long read assembly is much closer to delivering consistent complete plasmid assemblies than ever before, meant that both the right algorithms and the data were finally ready to make significant progress on plasmid analysis. The remaining difficulty was that, when software implementations for these models even exist, they require as input genomes that are already in the form of integer sequences. As the problem of how to choose markers for these sequences is not trivial, this is a significant barrier to using rearrangement distances in practice. We have implemented both integerisation (marker selection) and calculation of the DCJ-Indel distances within our tool pling, which finally makes the wider use of rearrangement distances possible. We also plan to further increase pling’s utility by allowing linear genomes as input, as ding is already able to handle them.

By using one subnetwork (with DCJ-Indel distances) embedded in another (with containment distances), pling generates a network whose topology also provides valuable information. The ability to interpret the network structure makes the reasons for a specific clustering clearer, which allows for better experimentation with thresholds, if a dataset requires it. Additionally, we use it to identify hubs, which connect many unrelated plasmids. Most of these are small (few kb) plasmids dominated by a transposon or IS element, but some are very large cointegrates which are presumably mosaics of many recombinations/fusions (data not shown). Simpler fusion events can also be identified – for example the presence of a cointegrate plasmid in Figure 4 is reflected in the network by it bridging the otherwise totally unrelated blue and orange clusters. This highlights the hierarchical and mosaic nature of the evolution of mobile elements. Importantly, fusion events are not rare – in one recent study 20% of plasmids in a database were found to be multireplicon [13], and in another it was found that 65% of plasmids from a historical collection (1917-1954) could be found completely embedded within modern descendants which were much larger [46].

Runtime for pling scales with the number of pairwise alignments, which depends on how many similar plasmids are in the dataset; the sourmash prefilter keeps the runtime relatively low by ensuring alignment is only done on similar genomes, with all datasets completing within two hours. RAM use on the other hand is well controlled (under 9Gb for all datasets), making computation for smaller datasets feasible on a laptop (in contrast to mge-cluster, which has high RAM use). Runtimes for mge-cluster and MOB-suite are generally faster for more similar datasets, but pling outperforms them on more divergent datasets.

On our empirical nosocomial datasets (Russian Dolls, Cambridge) we demonstrate that pling produces an interpretable network, which is in general more robust to structural changes and movement of TEs than MOB-suite or mge-cluster, and finds clusters with a larger core-genome (or backbone). Nevertheless, on the Russian Dolls dataset, MOB-suite (with secondary clustering) performs essentially identically to pling, successfully not clustering the pair of structurally diverged plasmids, because the mash distance is just above a threshold. On the other hand in the exemplar dataset (PSD) it would fail to separate the plasmids in Fig 3B, which have low enough mash distance to be clustered together with both primary and secondary MOB-suite clusterings. This highlights the benefit of the pling approach, which combines both containment/sharing information with DCJ-Indel/rearrangement information.

There are limitations to this study and approach. First, it is dependent on good genome assemblies, as errors in assembly will affect the DCJ-Indel distance. Our feeling is that as routine perfect bacterial genome assembly gradually becomes a reality via long reads, pling will become progressively more useful. Second, as with all network clustering approaches, cluster assignment is inevitably influenced by sampling. This is most apparent when trying to circumvent the influence of promiscuous TEs: we can only identify promiscuous transposons (for example) if they occur sufficiently often in the data. Nonetheless, even with default settings pling mediates this problem better than other clustering tools (e.g. note the detection of the Tn4401 transposon in the Russian Dolls data). We anticipate future work on developing other community detection algorithms which identify network topologies created by TEs. Third, we chose small datasets (Russian Dolls, N=62 and Addenbrookes, N=193) to compare clustering approaches to make manual curation of truth data possible, but mge-cluster is not meant to be used on datasets of less than a hundred (pers.comm Jukka Corander), and we did not run a full parameter sweep to explore if that improved clustering (as that would have required a lot of manual curation). Fourth, DCJ-Indel distance calculation is an NP-hard problem, which in theory can cause runtime problems. In practice, with ding’s highly efficient integer linear programming formulation, this has not been an issue, and the many pairwise alignments necessary for integerisation and containment distance calculation are typically the most resource heavy step. Finally, we did not benchmark against COPLA [18], a well known tool for analysing plasmids through Plasmid Taxonomic Units (PTUs). The first reason for this is that PTUs are intended to be wide groupings, much too broad to identify recent possible transmissions. The second is that PTUs are fixed, and COPLA simply assigns plasmids to predefined PTUs, rather than clustering plasmids, so we believe the methods are not comparable.

We have shown, both through constructed and real hospital datasets, that this method produces plasmid clusters with a recent common evolutionary history, removing false associations between plasmids. This greatly reduces the need for manual curation of plasmid clusters, potentially significantly simplifying epidemiological studies. Furthermore, we believe this concept of relatedness will be of utility in future evolutionary work, as DCJ-Indel represents the number of structural genetic events distinguishing two plasmids. We envisage revisiting previous plasmid network-based analyses through this lens [47, 48].

## Author statements

### Author contributions

Conceptualisation: DF, ZI

Data Curation: DF

Formal analysis: DF, LR

Funding acquisition: JS, ZI

Investigation: DF

Methodology: DF, LL, LR, LB, RW, JS, ZI

Project administration: ZI

Resources: LR, ZI

Software: DF, LL, LB

Supervision: ZI

Validation: DF

Visualisation: DF, LB

Writing – original draft: DF

Writing – review & editing: DF, LL, LR, LB, RW, ZI

### Conflicts of interest

The authors declare that there are no conflicts of interest.

### Funding Information

DF received funding from the European Union’s Horizon 2020 research and innovation programme under Marie SkŁodowska-Curie grant agreement No 956229 (ALPACA).

## Acknowledgements

We would like to thank Martin Hunt, Sally Partridge and Ed Feil for discussions and advice, and Marjorie Gibbon, Natacha Couto and Vesa Quarkaxhija for testing and evaluating pling. We would also like to thank Jukka Corander and Anita Schürch for discussions and advice on applicability of mge-cluster.

## Notes

https://github.com/iqbal-lab-org/pling

https://github.com/babayagaofficial/pling_paper_analyses

### Competing Interest Statement

The authors have declared no competing interest.

