## Supplementary information and figures for "Applying rearrangement distances to enable plasmid epidemiology with pling"

*Programs and parameters used for alignment in pling*

To integerise from alignment, we use MUMmer 3 [30], specifically the nucleotide alignment subprogram nucmer with parameters diagdiff=20, breaklen=500, and flag maxmatch. Then we use delta-filter with the flag -1 to filter out only the "best" matches – so if two matches map the same interval, only the one with the better score is kept. If the two matches have the same score, both are kept. We also use show-coords and show-snps to format the nucmer output and generate SNPs. We filter out matches whose identity is less than 80% in the show-coords step.

*Error handling in integerisation from alignment in pling*

When projection between coordinates is unreliable, two errors can occur while resolving overlaps:

1. The ordering of the matches is broken
2. A match where start coordinate is greater than end coordinate is created

To catch error one, the algorithm checks that the order is maintained when splitting matches, and also checks every five iterations that the order has not been broken. In the first case, the overlap fixing is broken off and the previous version of matches is used for assigning integers. In the second, the version from five iterations ago is used. To catch the second type of error, anytime a new match is created, the algorithm checks that its coordinates are valid. If not, again overlap fixing is broken off and the previous version of matches is used for assigning.

Breaking off overlap fixing prematurely means that we end up not using fully resolved matches when assigning integers, which means information about the presence of duplicate regions can be missing from the integerisation. This then affects the DCJ-Indel distance calculation, and can result in distances that are slightly off. This is not generally an issue for the clustering though, as the edge cases that necessitate breaking off overlap fixing are typically very divergent plasmids, and a slight difference in distance would not result in them gaining a DCJ-Indel edge in the final network. Therefore the approximate integerisation, which results in an approximate DCJ-Indel distance, is sufficient for pling.

*Gene annotation workflow*

Similar to the workflow that implements integerisation from alignment, we start by building a containment network, but for the annotation workflow this step does not also produce integer sequence representations of the genomes. We then annotate each plasmid using bakta [31], followed by running panaroo [32] on each containment community. Then for each community, using the pangenome outputted by panaroo, we reconstruct the order of genes in each plasmid by mapping each gene back to the plasmid with minimap2 [49], from which we can then construct integer sequences. These integer sequences will have the same integer label assigned to each gene across the whole plasmid community. We then simplify the integer sequences by finding blocks of genes in conserved order across all the plasmids in the community, and representing these blocks with a single integer. This gives us our final integer representation, which we use for the rest of the workflow, which is the same as previously described.

*Running clustering tools and sourmash*

We clustered the datasets with the version of pling at commit https://github.com/iqbal-lab-org/pling/commit/ed8ba1e84bbffad9963d71aa2db1c43bd67effaf. Note that later development mostly involved refactoring and bug fixes that did not affect functionality, but running with a different version may produce a permutation of cluster labels. For mge-cluster (v1.1.0), we used the --create flag with the min_cluster parameter set to 2, as we wanted to allow pairs of plasmids to be considered a cluster. All other parameters were run with defaults. For MOB-suite (v3.1.8), we ran subcommand mob_typer with the --multi flag to generate the necessary input file to then run subcommand mob_cluster in build mode, which is the clustering subcommand. We used MOB-suite default parameters. Similarly, we ran mash [41], and sourmash [21] with --max-containment flag, using defaults.

*Runtime evaluations*

We ran runtime evaluations on pling v1.0.3. We constructed four datasets to evaluate runtimes: 300 IncF, 1500 IncF, 300 type blind and 1500 type blind. To construct the 1500 type blind dataset, we sampled 1500 plasmids from PLSDB, and for the 300 type blind further subsampled from the set of 1500. These are both sets of truly random plasmids, which would have a high degree of diversity. We also chose to create datasets from sampling just IncF plasmids from PLSDB – the intention was to create a dataset where plasmids had a higher degree of similarity, while still maintaining some diversity. We ran the size 300 and size 1500 datasets under different scenarios: for the datasets of size 300, we ran pling on 8 threads on a laptop with 8 cores and 16GB memory; for the datasets of size 1500, we ran on a server with 48 threads and recorded maximal memory usage. We wanted to do equivalent comparisons for MOB-suite and mge-cluster, but were hampered by installation problems. MOB-suite we were only able to install on the laptop, and not on the compute cluster that we used for the larger datasets, whereas for mge-cluster, we were able to install it on the compute cluster but not on the laptop. As a result we were unable to evaluate MOB-suite for the larger sized datasets; for mge-cluster we imitated the laptop evaluation by allocating 8 cores and 16GB memory on the compute cluster. Mge-cluster failed the laptop test, as its RAM requirements surpassed 16GB. The remaining results are recorded in Supplementary Table 1.

*Alignment visualisations*

We used a visualisation script by Martin Hunt (https://github.com/martinghunt/bioinf-scripts/blob/master/python/multi_act_cartoon.py), which runs nucmer and then creates figures from the output.

*Analysis of Addenbrookes hospital plasmids*

To assess how well each tool's clustering corresponded to the Addenbrookes data "truth", we evaluated the minimum number of plasmids that would need to be reassigned to get from the tool's clustering to the truth clustering. This is equivalent to the distance introduced by

van Dongen [44–46]. The results were 12 for pling, 23 for MOB-suite and 81 for mge-cluster. We implemented a python script to calculate these distances on the basis of equation 12 in [45]. We get the percentage of plasmids that require reassignment by dividing by the total number of plasmids (N=193). Note that we are able to interpret this number as the percentage of plasmids requiring reassignment due to knowing that there are no disagreements arising due to plasmids being split off a truth cluster and then assigned to a different truth cluster – in the case of such an event the van Dongen distance would count each reassigned plasmid twice, making the interpretation of the division of the van Dongen distance by the number of data points more ambiguous.
We determined core genes (i.e. genes present in each genome) per pling cluster on the Addenbrookes hospital dataset by running ggcaller (v1.3.4) [50] with default parameters and the --repeat flag. We summed the length of the core genes to get the core genome size, and then divided by genome length to get the relative core genome size per plasmid. We chose the median to represent the average relative core genome size within a cluster, so that size outliers do not affect the value excessively. We used python library seaborn [51] for data visualisation.
Phylogenies were built from E. coli and K. pneumoniae complete chromosomes [43] using Parsnp (v1.7.4) [52] at default settings with a randomly selected internal reference. The trees are visualised with the help of python library ete3 [53].

### Supplementary Figures

| tool | dataset | sourmash | Gurobi | run time | max memory | cores |
|---|---|---|---|---|---|---|
| pling | 300 IncF | yes | yes | 22 minutes 18 seconds | - | 8 |
| pling | 300 IncF | yes | no | 24 minutes 7 seconds | - | 8 |
| MOB-suite | 300 IncF | - | - | 4 minutes 30 seconds | - | 8 |
| pling | 300 type blind | yes | yes | 2 minutes 10 seconds | - | 8 |
| pling | 300 type blind | yes | no | 2 minutes 54 seconds | - | 8 |
| MOB-suite | 300 type blind | - | - | 3 minutes 14 seconds | - | 8 |
| pling | 1500 IncF | yes | yes | 1 hour 19 minutes 19 seconds | 6.67G | 48 |
| pling | 1500 IncF | no | yes | 4 hours 33 minutes 19 seconds | 8.6G | 48 |
| pling | 1500 IncF | yes | no | 1 hour 54 minutes 42 seconds | 6.69G | 48 |

| | | | | | | |
|---|---|---|---|---|---|---|
| mge-cluster | 1500 IncF | - | - | 3 minutes 16 seconds | 84.93G | 48 |
| pling | 1500 type blind | yes | yes | 6 minutes 38 seconds | 6.13G | 48 |
| pling | 1500 type blind | no | yes | 1 hour 44 minutes 15 seconds | 8.61G | 48 |
| pling | 1500 type blind | yes | no | 10 minutes 26 seconds | 6.24G | 48 |
| mge-cluster | 1500 type blind | - | - | 24 minutes 24 seconds | 47.61G | 48 |

**Supp. Table 1 Runtime comparisons between pling, mge-cluster and MOB-suite.** We evaluate on four datasets: 300 randomly selected plasmids from PLSDB (300 type blind), 1500 randomly selected plasmids from PLSDB (1500 type blind), 300 randomly selected plasmids from PLSDB that contain an IncF replicon (300 IncF), and 1500 randomly selected plasmids from PLSDB that contain an IncF replicon (1500 IncF). We run pling with and without sourmash or Gurobi to demonstrate runtime improvements, and also compare to mge-cluster and MOB-suite where possible. Tests were done for two scenarios: smaller dataset on an 8 core laptop with 16GB memory, and larger dataset on a 48 core server. For the smaller datasets we just measure runtimes, for the larger datasets we also evaluate memory usage.

| Example | Plasmid A | Plasmid B | Jaccard | Containment | DCJ-Indel | Length A | Length B |
|---|---|---|---|---|---|---|---|
| A | pKPC_CAV1321-45 | pKPC_CAV1668 | 0.00108342 | 0 | 1 | 44,846 | 43,433 |
| B | NZ_AP024098.1 | NZ_CP022793.1 | 0.00473708 | 0.07220216606 | 88 | 2139380 | 1989056 |
| C | cpe027_3 | cpe060_4 | 0.0909306 | 0 | 1 | 115672 | 8514 |
| D | pKPC_CAV1320 | pKPC_CAV1043 | 0.0567566 | 0.2 | 2 | 13,981 | 59,138 |
| D | pKPC_CAV1320 | pKPC_CAV1492 | 0.0621044 | 0.2 | 2 | 13981 | 69,158 |
| D | pKPC_CAV1320 | pKPC_CAV1335 | 0.0843302 | 0.2 | 2 | 13981 | 113,105 |
| D | pKPC_CAV1320 | pKPC_CAV1596-97 | 0.0766828 | 0.2 | 2 | 13981 | 96,702 |
| D | pKPC_CAV1320 | pKPC_CAV1321-45 | 0.0480354 | 0.2 | 2 | 13981 | 44,846 |

**Supp. Table 2 Genetic distances between pairs from PSD.** Jaccard (mash, non-shared kmers over union), containment (sourmash, kmers unique to smaller plasmid over kmers in smaller plasmid), and DCJ-Indel distances between pairs of plasmids in Figure 3. Note that for pairs in example D of Figure 3, we only show the distances between the first plasmid (i.e. the plasmid dominated by a promiscuous TE), and all other plasmids.

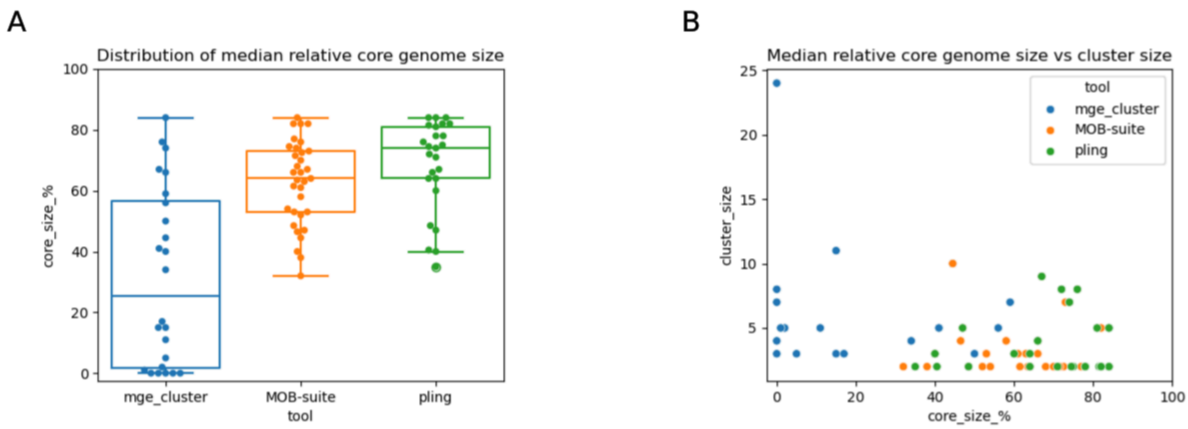


**Supp. Fig. 1 Core genome sizes and clusters.** A) Box plot of (median) core genome size as a percentage of genome size, for a cluster B) Core genome size as a percentage of genome size (median value per cluster) plotted against cluster size, for each tool tested.

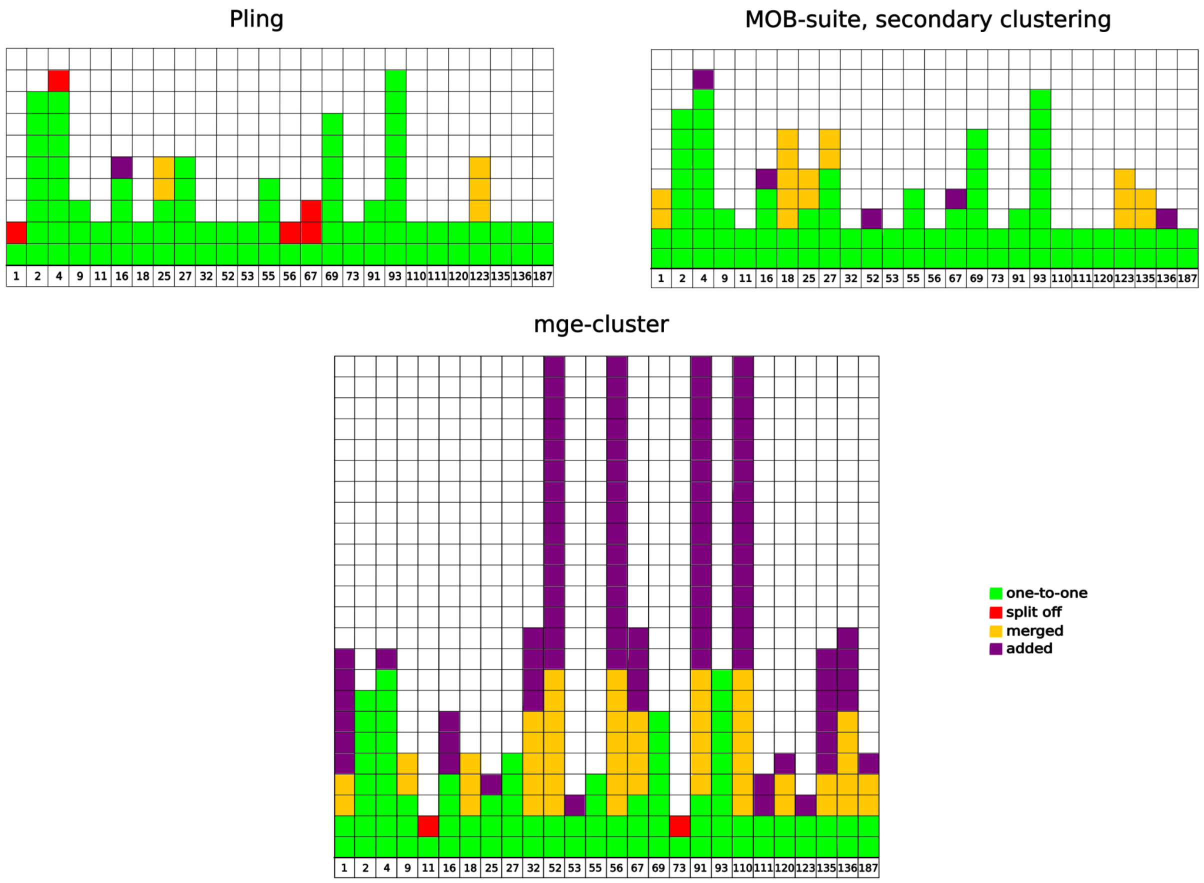


**Supp. Fig. 2 Representation of plasmid assignments in the "truth" clustering vs in the tool clustering.** A single block represents one plasmid; a column is one truth cluster, with labelling at the bottom. Green blocks mean a plasmid was mapped to the same cluster as in the truth; red blocks indicate the plasmid was split off from the "truth" cluster. Yellow blocks show plasmids added to the "truth" cluster due to merging of clusters; purple blocks signify the addition of a singleton plasmid to a "truth" cluster. Note that merges and additions are counted multiple times in this visualisation – for example, pling merges clusters 25 and 123, and there are yellow blocks in both columns for these clusters, but these signify the same merge event.

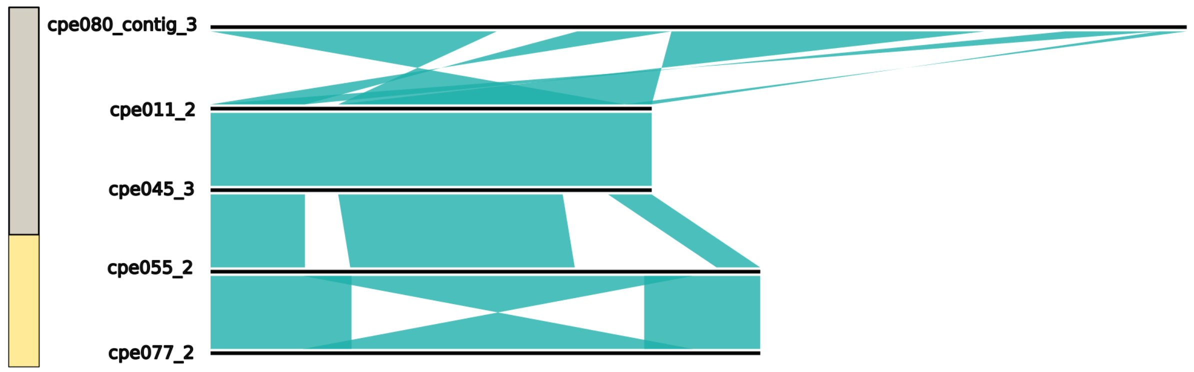


**Supp. Fig. 3 Merged "truth" clusters 25 and 123 alignment.** An alignment of clusters 25 and 123, which are merged into a single cluster by pling. The coloured bars correspond to the cluster assignment in the truth, and shaded areas between adjacent plasmids indicate shared regions. As we can see from the alignments, all these plasmids share a backbone.

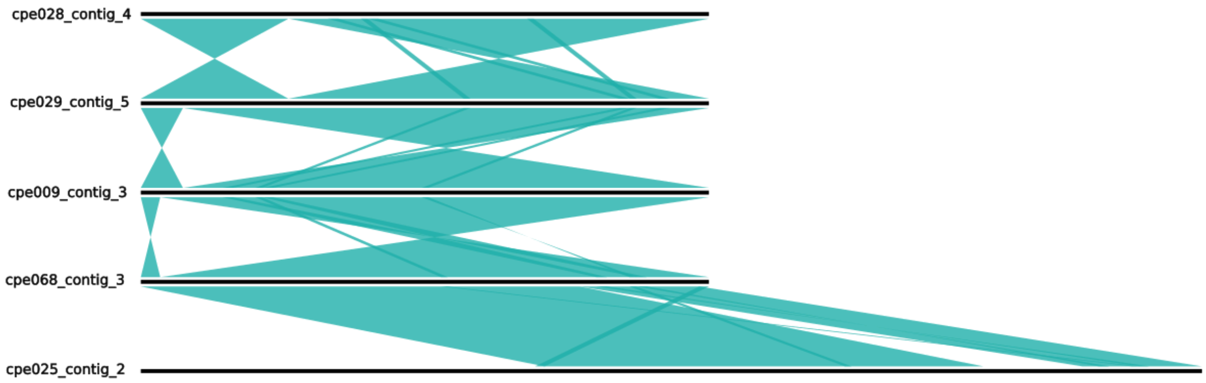


**Supp. Fig. 4 "Truth" cluster 16 with singleton.** An alignment of cluster 16 and the singleton which pling adds. The singleton is the bottommost plasmid. Shaded areas between adjacent plasmids indicate shared regions. The added singleton fully contains the smaller plasmids from cluster 16, which makes their close relationship apparent.

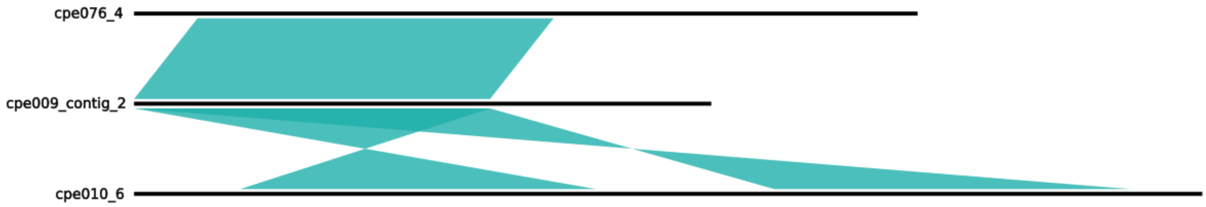


**Supp. Fig. 5 Novel cluster of three.** An alignment of a novel cluster pling finds. Shaded areas between adjacent plasmids indicate shared regions. All three plasmids share a backbone that has undergone no rearrangements, only a duplication in the largest plasmid.

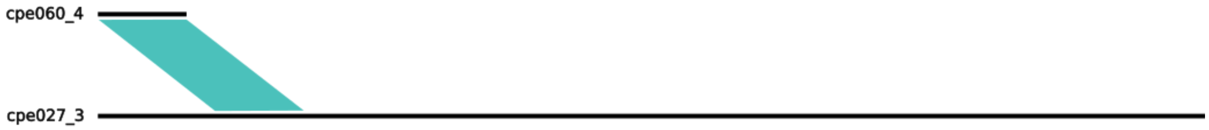

**Supp. Fig. 6 Novel pair 1.** An alignment of a novel pair pling finds. Shaded areas between adjacent plasmids indicate shared regions. The larger plasmid shares a replicon (ColRNAI_rep_cluster_1987) with the smaller, and also contains four additional replicons (IncFIA, IncFIB, IncFIC, and IncFII), indicating this is likely a cointegrate plasmid and one of its parents.

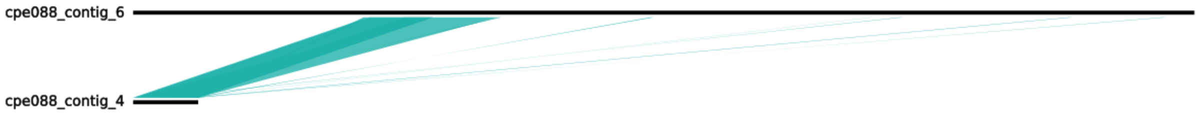


**Supp. Fig. 7 Novel pair 2.** An alignment of a novel pair pling finds. Shaded areas between adjacent plasmids indicate shared regions. The smaller plasmid is fully contained in the larger, and duplicated. Both plasmids are rep type IncFIB, and from the same isolate.

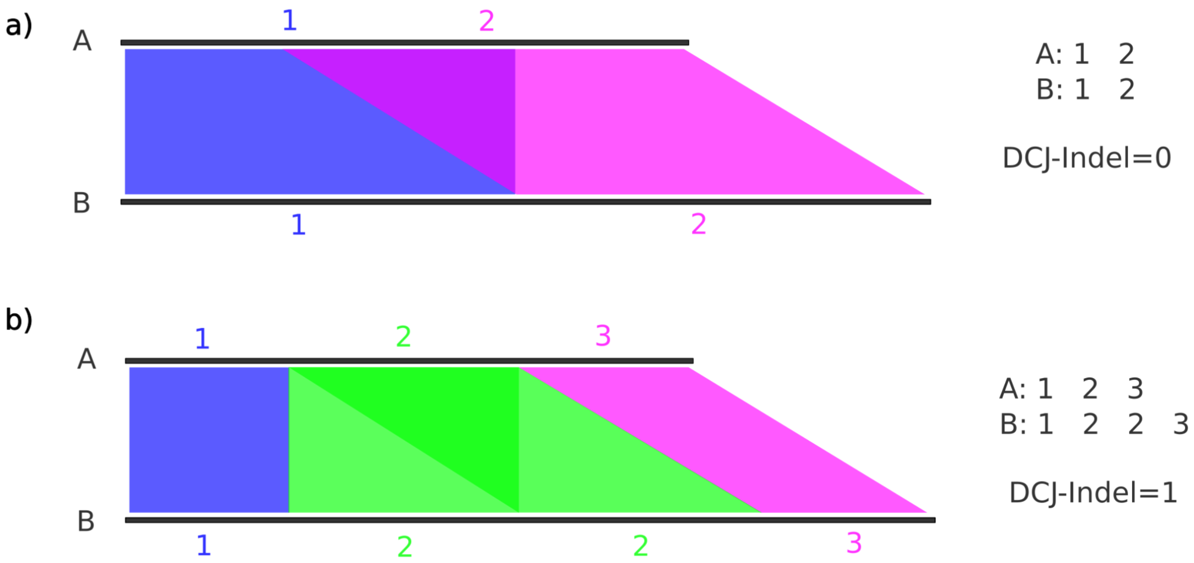


**Supp. Fig. 8 Example of how not accounting for overlaps results in incorrect DCJ-Indel distances.** In panel a), no attention is paid to whether or not there are overlaps, so genomes A and B are given the same integerisation ("1 2"), which would lead a user to believe they were structurally the same (with DCJ-Indel distance of 0), when they are not. In panel b) the overlapping matches between A and B are noticed, allowing us to identify that the green region labelled 2 is seen twice in genome B.

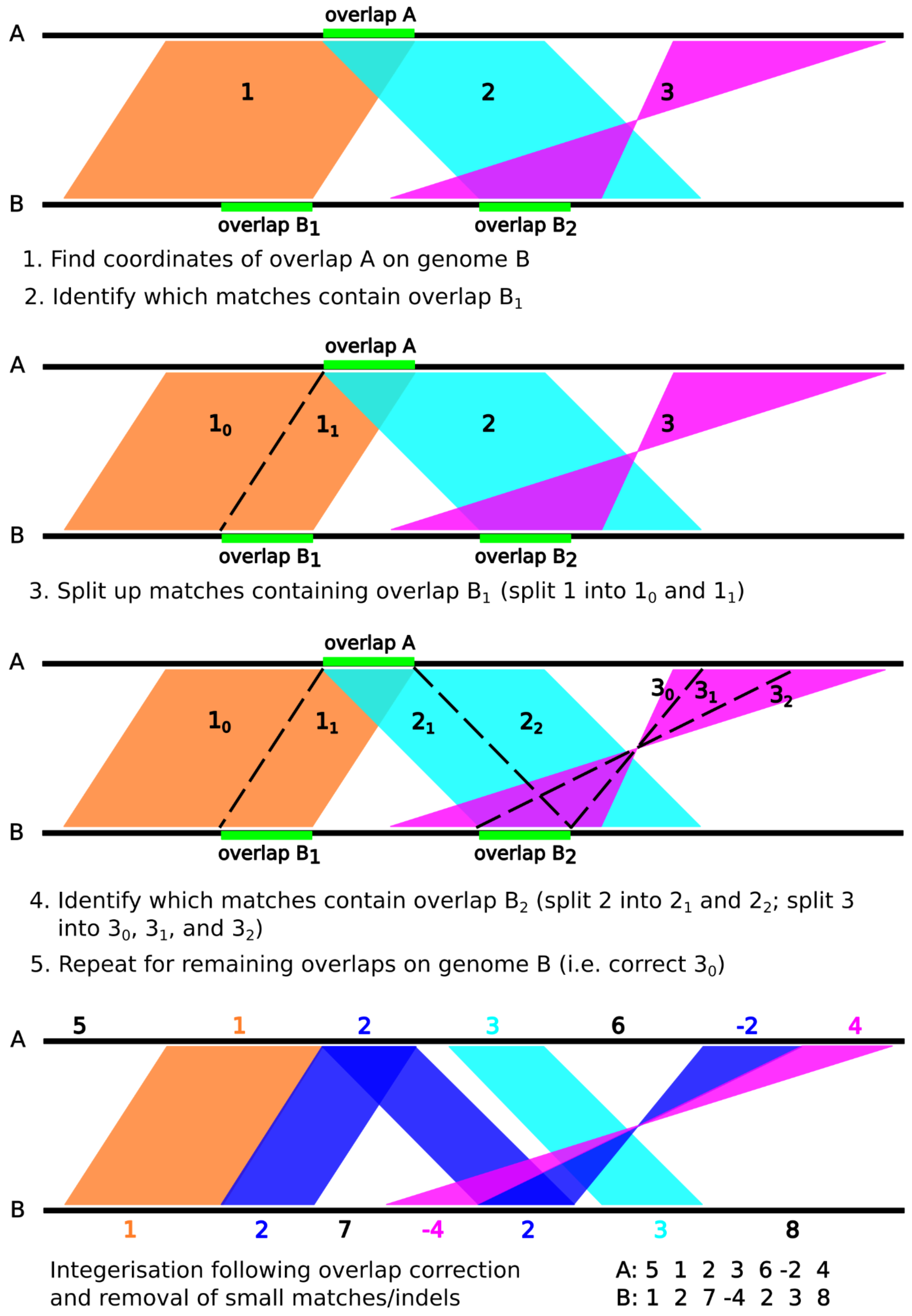


**Supp. Fig. 9 Schematic showing how overlap fixing works.**
